# N1 cassette-lacking NMDA receptors mediate the antidepressant activity of ketamine

**DOI:** 10.1101/2025.04.12.648536

**Authors:** Alina T. He, Wenbo Zhang, Hongbin Li, YuShan Tu, Dongju Lee, Doyeon Kim, Zhengping Jia, Lu-Yang Wang, Michael W. Salter

## Abstract

Ketamine has emerged as a rapid-acting and robust antidepressant^1,2^. However, the mechanism of its antidepressant action remains enigmatic. The core issue that has yet to be resolved is whether NMDA receptors (NMDARs), which are subject to open channel blockade by ketamine^3,4^, mediate the antidepressant effect. NMDARs naturally undergo alternative splicing of the obligatory GluN1 subunit^5^, producing receptor diversity in the brain that has not been considered in the actions of ketamine. Here we discover that alternative splicing of *Grin1* exon 5, which leads to exclusion (GluN1a) or inclusion (GluN1b) of the N1 cassette, located in the N-terminal domain of GluN1 distant from the pore, unexpectedly dictates the level and dynamics of NMDAR blockade by ketamine and gates its antidepressant activity. We find that ketamine prevents NMDAR-dependent long-term potentiation (LTP) in the CA1 region of the hippocampus in mice engineered to exclude *Grin1* exon 5 (GluN1a mice), but ketamine has no effect on LTP in mice engineered to include this exon (GluN1b mice). Ketamine inhibits synaptic NMDARs in CA1 pyramidal neurons in both GluN1a and GluN1b mice, with the level of steady-state blockade marginally greater in GluN1a- than in GluN1b-containing NMDARs. However, the rate of relief of ketamine blockade upon membrane depolarization is markedly slower in GluN1a than in GluN1b neurons such that GluN1a-containing receptors remain blocked during bursting activity, whereas those containing GluN1b escape the ketamine blockade. Furthermore, ketamine treatment, either via systemic administration or local infusion into the hippocampus, induces an antidepressant effect in GluN1a mice but has no effect in GluN1b mice. Collectively, we identify GluN1a-containing NMDARs, which are persistently blocked by ketamine during neuronal firing activity, to be selectively responsible for the antidepressant effect.

## Introduction

Major depression is a prevalent and debilitating disorder, affecting hundreds of millions of people worldwide and causing tremendous personal and socioeconomic burdens^6,7^. Conventional antidepressants typically require weeks to act and are ineffective in around one-third of patients^8,9^. In contrast, ketamine alleviates depressive mood within hours and is effective even in patients resistant to traditional antidepressants^1,2^. Although the treatment of depression has been revolutionized by the clinical use of ketamine, the primary target mediating the rapid onset and highly effective antidepressant action has yet to be resolved and remains a topic of intense controversy.

The debate has focused on whether the antidepressant effect of ketamine is mediated through its blockade of NMDARs or through mechanisms independent of these receptors^10–16^. NMDARs are remarkably diverse in their structural and functional properties, depending on their subunit composition^17^. These receptors are heterotetrameric assemblies composed of two obligatory GluN1 subunits and typically two GluN2 subunits that bind the co-agonists glycine and glutamate, respectively. Variation among the four GluN2(A-D) subunits is commonly regarded as the determinant of diversity in NMDAR function and pharmacology^17^. However, a greater number of subtypes exist for GluN1, with eight isoforms generated by alternative splicing of *GRIN1*^5^, the functional consequences of which have been largely overlooked.

*GRIN1* alternative splicing yields isoforms excluding exon 5 (GluN1a) or including this exon (GluN1b), which encodes the N1 cassette, a highly evolutionarily conserved 21-amino acid sequence^5,18^. GluN1 isoforms lacking or containing the N1 cassette are normally expressed in the adult brain across species^18–21^. The N1 cassette is located in the extracellular N-terminal domain at an interface between GluN1 and GluN2 ligand binding domains^22^. Here we investigated whether the presence of the N1 cassette has consequences for the ketamine blockade of NMDARs and its antidepressant activity.

### Ketamine inhibition of LTP is prevented by inclusion of the N1 cassette

To investigate whether the N1 cassette alters the effects of ketamine, we used mice engineered to exclude *Grin1* exon 5 (GluN1a mice) or include this exon (GluN1b mice) to examine a well-established function of NMDARs, inducing long-term potentiation (LTP) at Schaffer collateral-CA1 synapses of the hippocampus^23,24^. GluN1a and GluN1b mice have been characterized and do not differ from wild-type (WT) mice in total *Grin1* mRNA expression or in basal synaptic transmission, NMDAR current amplitudes, or NMDA to AMPA ratio at Schaffer collateral-CA1 synapses^18^. We recorded extracellular field excitatory postsynaptic potentials (fEPSPs) in acute hippocampal slices from GluN1a and GluN1b mice, as well as their WT littermates. After establishing a stable baseline of fEPSPs for at least 20 min, slices were treated with ketamine (10 μM) or untreated in controls (Fig. 1). In all genotypes, fEPSP slopes 1 h later in ketamine-treated slices were not different from those in untreated controls (Extended Data Fig. 1A). We next applied theta-burst stimulation (TBS: 15 bursts at 5 Hz, with each burst comprised of 4 stimuli at 100 Hz) to induce LTP. This stimulation pattern mimics theta frequency firing observed in hippocampal pyramidal neurons in awake behaving animals^25–27^. In WT slices, the magnitude of LTP in ketamine-treated slices was significantly less than that in control slices (control = 140 ± 1.9%, ketamine = 118 ± 2.4%, p = 0.0003; Fig. 1A, B and Extended Data Fig. 1B), consistent with previous reports of ketamine inhibition of LTP^28–30^. In addition, we found that in GluN1a slices, LTP magnitude was significantly less in ketamine-treated slices than that in untreated controls (control = 147 ± 5.4%, ketamine = 112 ± 3.5%, p < 0.0001; Fig. 1C, D and Extended Data Fig. 1B). In contrast, LTP magnitude was not different in GluN1b slices with or without ketamine treatment (control = 147 ± 5.6%, ketamine = 140 ± 5.8%, p = 0.287; Fig. 1E, F and Extended Data Fig. 1B). Furthermore, LTP following ketamine treatment was significantly less in WT or GluN1a slices compared to in GluN1b slices (p = 0.0008, p < 0.0001, respectively; Fig. 1 and Extended Data Fig. 1B). Therefore, we conclude that inclusion of the N1 cassette in GluN1 prevents ketamine inhibition of LTP.

**Figure 1.**
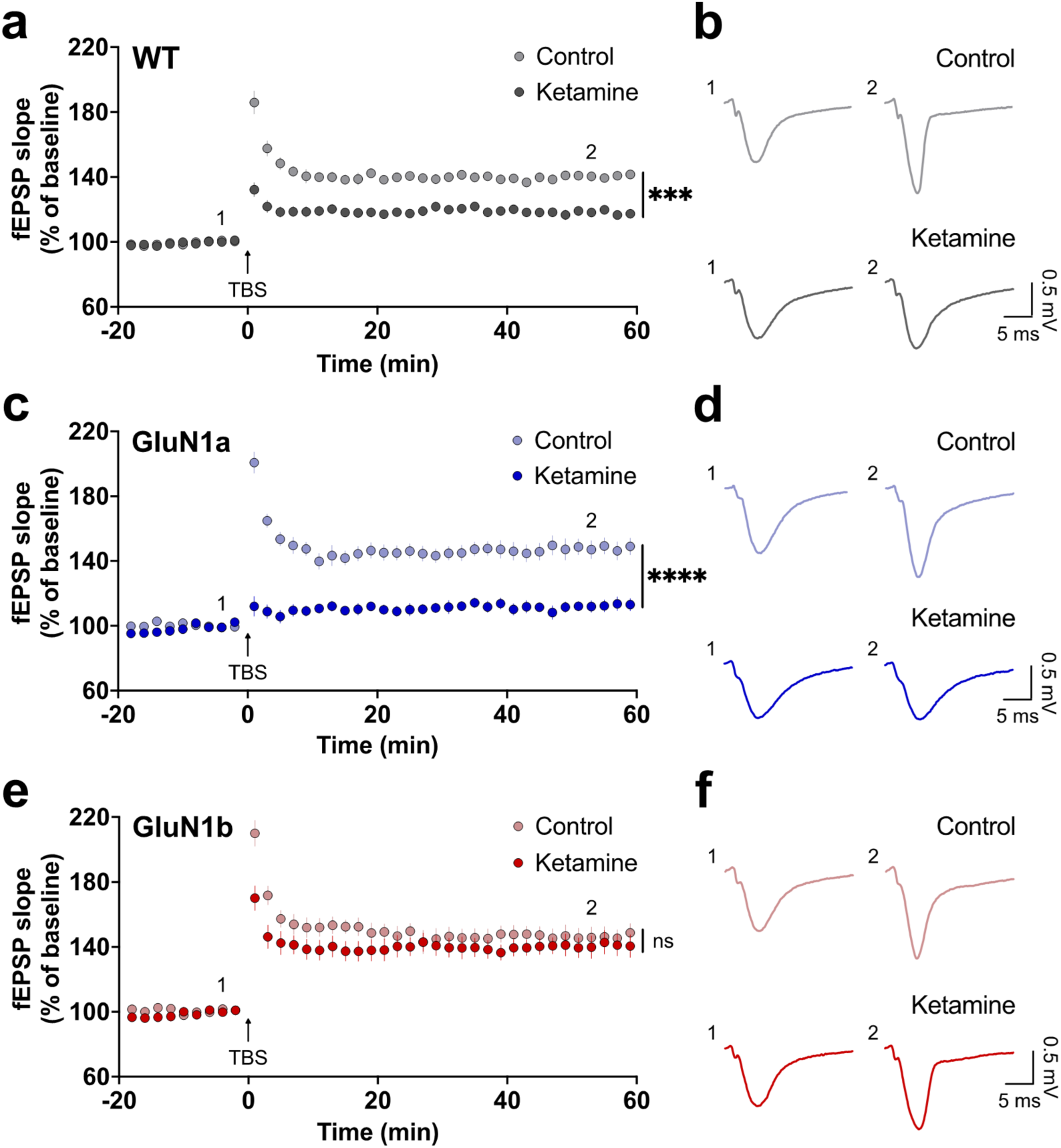
Ketamine inhibits LTP at Schaffer collateral-CA1 synapses in GluN1a but not in GluN1b hippocampal slices. Left, time plots of fEPSP slopes from **(a)** WT (control n = 9, ketamine n = 13), **(c)** GluN1a (control n = 9, ketamine n = 9), and **(e)** GluN1b (control n = 8, ketamine n = 9) mice. Slices were treated with ketamine (10 µM) for 1 h or untreated in controls. TBS was delivered at time 0. fEPSP slopes are normalized to the 10 min before TBS. Right, representative fEPSP traces from time points 1 and 2 from **(b)** WT, **(d)** GluN1a, and **(f)** GluN1b mice. Data are represented as mean ± SEM. ***p < 0.001, ****p < 0.0001, ns p > 0.05, two-way ANOVA with Tukey’s post hoc test.

### Inclusion of the N1 cassette attenuates ketamine block of NMDARs

Given that ketamine suppressed LTP in GluN1a mice but had no effect on LTP in GluN1b mice, we hypothesized that ketamine may differentially inhibit GluN1a-versus GluN1b-containing NMDARs. We examined the effects of ketamine on synaptic NMDAR currents in GluN1a, GluN1b, and WT mice. In whole-cell patch-clamp recordings from CA1 pyramidal neurons in hippocampal slices, NMDAR excitatory postsynaptic currents (EPSCs) were evoked by Schaffer collateral stimulation and pharmacologically isolated using NBQX (10 μM) to block AMPA receptors and bicuculline (10 μM) to block GABA_A_ receptors. The membrane potential was held at −40 mV to partially relieve the voltage-dependent Mg^2+^ blockade of NMDARs. After establishing a stable baseline of at least 10 min, slices were treated with ketamine (10 μM) or untreated in controls (Fig. 2A, B). NMDAR EPSC amplitudes after 30 min were significantly reduced with ketamine treatment in all genotypes (WT = 44.1 ± 2.7%, GluN1a = 38.3 ± 3.9%, GluN1b = 56.7 ± 5.3% of baseline, p < 0.0001; Fig. 2A, B). This decrease in NMDAR EPSC amplitudes in GluN1a neurons was significantly greater than that in GluN1b neurons (p = 0.0219; Fig. 2A, B). In addition, ketamine had no effect on NMDAR EPSC rise or decay times in any genotype (Extended data Fig. 2).

**Figure 2.**
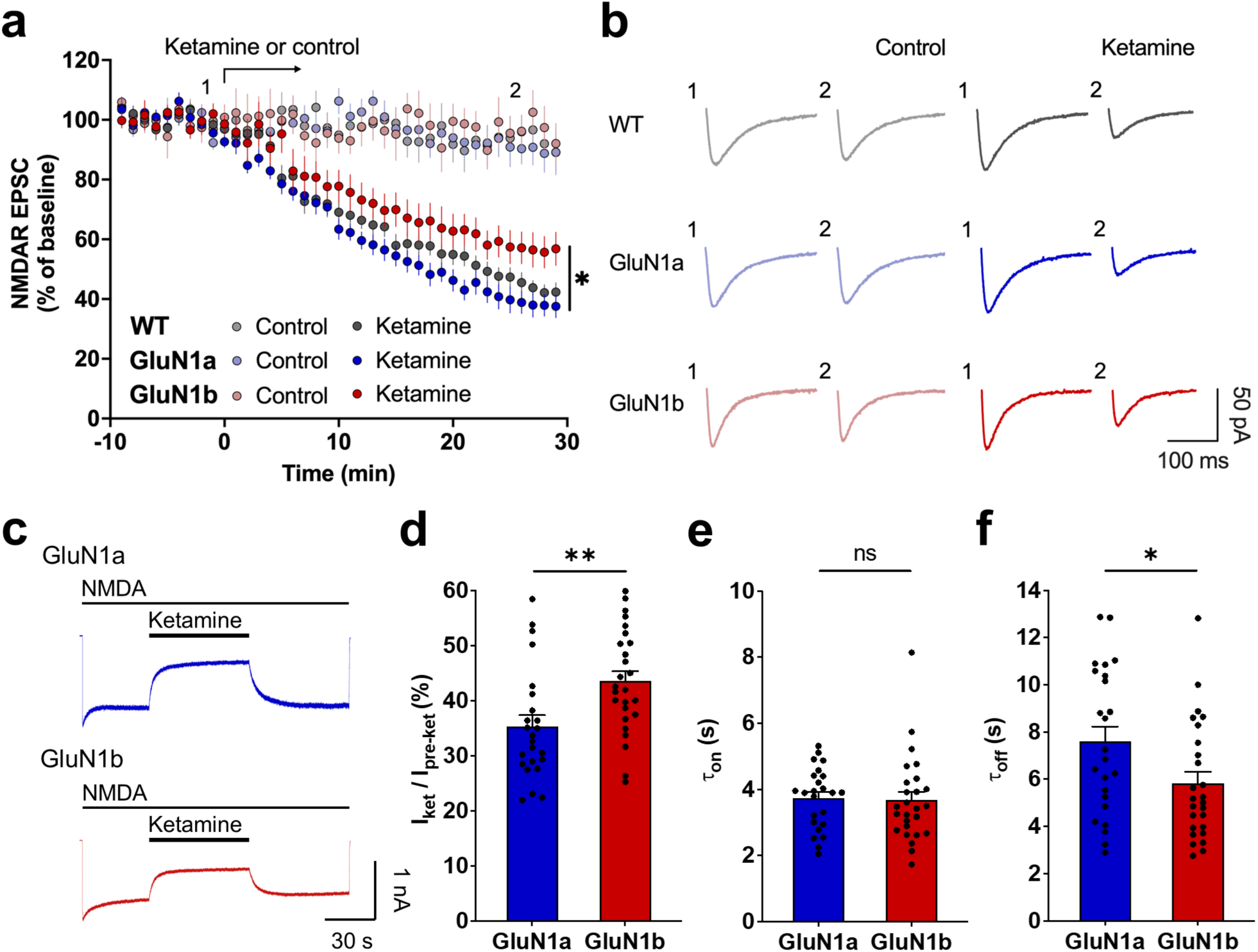
Ketamine inhibition is greater in GluN1a- than in GluN1b-containing NMDARs. **a,** Time plot of NMDAR EPSC amplitudes from CA1 pyramidal neurons holding at −40 mV in hippocampal slices from WT (control n = 7, ketamine n = 8), GluN1a (control n = 8, ketamine n = 8), and GluN1b (control n = 7, ketamine n = 8) mice. Slices were treated with ketamine (10 μM) for 30 min starting at time 0 or untreated in controls. NMDAR EPSC amplitudes are normalized to the baseline. **b,** Representative NMDAR EPSC traces at time points 1 and 2. Data are represented as mean ± SEM. *p < 0.05, two-way ANOVA with Tukey’s post hoc test. **c,** Representative traces of NMDA-evoked currents in hippocampal neurons cultured from GluN1a (n = 24) and GluN1b (n = 26) mice. The membrane potential was held at −40 mV. Ketamine (10 μM) was applied for 1 min during NMDA administration. **d,** NMDA-evoked current amplitudes during ketamine treatment relative to that before ketamine application. Tau on **(e)** and Tau off **(f)** of ketamine. Data are represented as mean ± SEM. *p < 0.05, **p < 0.01, ns p > 0.05, Student’s t-test or Mann-Whitney test, as appropriate.

We examined the rate of ketamine inhibition of NMDAR currents using fast application of ketamine (10 μM) during administration of NMDA (50 μM plus glycine 1 μM) in whole-cell voltage-clamp recordings from hippocampal neurons cultured from GluN1a and GluN1b mice (Fig. 2C-F). The membrane potential was held at −40 mV, as neurons were bathed in Mg^2+^-containing extracellular solution. Similar to the NMDAR EPSCs, the reduction of NMDA-evoked currents by ketamine in GluN1a neurons was greater than that in GluN1b neurons (GluN1a = 35.3 ± 2.0%, GluN1b = 43.5 ± 1.8% of pre-ketamine current, p = 0.0044; Fig. 2D). We found that the rate of onset of ketamine inhibition was not different between GluN1a and GluN1b neurons (GluN1a τ = 3.75 ± 0.19 s, GluN1b τ = 3.67 ± 0.26 s, p = 0.825; Fig. 2E). However, the rate of recovery upon ketamine washout was significantly slower in GluN1a neurons compared to that in GluN1b neurons (GluN1a τ = 7.60 ± 0.63 s, GluN1b τ = 5.82 ± 0.49 s, p = 0.0310; Fig. 2F). Altogether, ketamine preferentially inhibited both synaptic NMDAR currents and NMDA-evoked currents in GluN1a neurons over GluN1b neurons.

### Inclusion of the N1 cassette accelerates voltage-dependent relief of ketamine blockade

An apparent paradox in our findings was that while ketamine abolished LTP in GluN1a slices and had no effect on LTP in GluN1b slices, there was only a modest difference in the level of steady-state inhibition of NMDAR currents between GluN1a and GluN1b neurons. We noted that ketamine inhibition was measured while holding the membrane at a constant negative potential, but during TBS induction of LTP the membrane potential repeatedly depolarizes with each burst of stimuli^31–33^. Membrane depolarization has been shown to relieve ketamine blockade from NMDARs^3^. We thus investigated the possibility that this voltage-dependent unblocking is different between GluN1a- and GluN1b-containing NMDARs. In recordings from hippocampal cultured neurons, we allowed ketamine inhibition of NMDA-evoked current to develop in Mg^2+^-free extracellular solution. After ketamine inhibition stabilized, we stepped the membrane potential from −60 mV to +60 mV to induce relief of blockade during continued ketamine application (Fig. 3A). In both GluN1a and GluN1b neurons, voltage-dependent unblocking was observed as a progressively increasing outward current upon stepping to +60 mV (Fig. 3B), which did not occur in the absence of ketamine (Extended data Fig. 3). We found that the maximum current at +60 mV was significantly less in GluN1a neurons than that in GluN1b neurons (GluN1a = 0.573 ± 0.083, GluN1b = 0.755 ± 0.057 of pre-ketamine current, p = 0.0387; Fig. 3D). Moreover, the rate of voltage-dependent relief of ketamine inhibition was 2.6-fold slower in GluN1a neurons compared to that in GluN1b neurons (GluN1a τ = 1.46 ± 0.30 s, GluN1b τ = 0.571 ± 0.19 s, p = 0.0075; Fig. 3C). Note that a difference in the level of ketamine inhibition between GluN1a and GluN1b neurons was dependent upon the presence of extracellular Mg^2+^ (Fig. 3A, Fig. 2D, Extended data Fig. 4). The substantially slower rate of voltage-dependent relief of inhibition in GluN1a than in GluN1b neurons was also observed in the presence of extracellular Mg^2+^ (Extended data Fig. 4).

**Figure 3.**
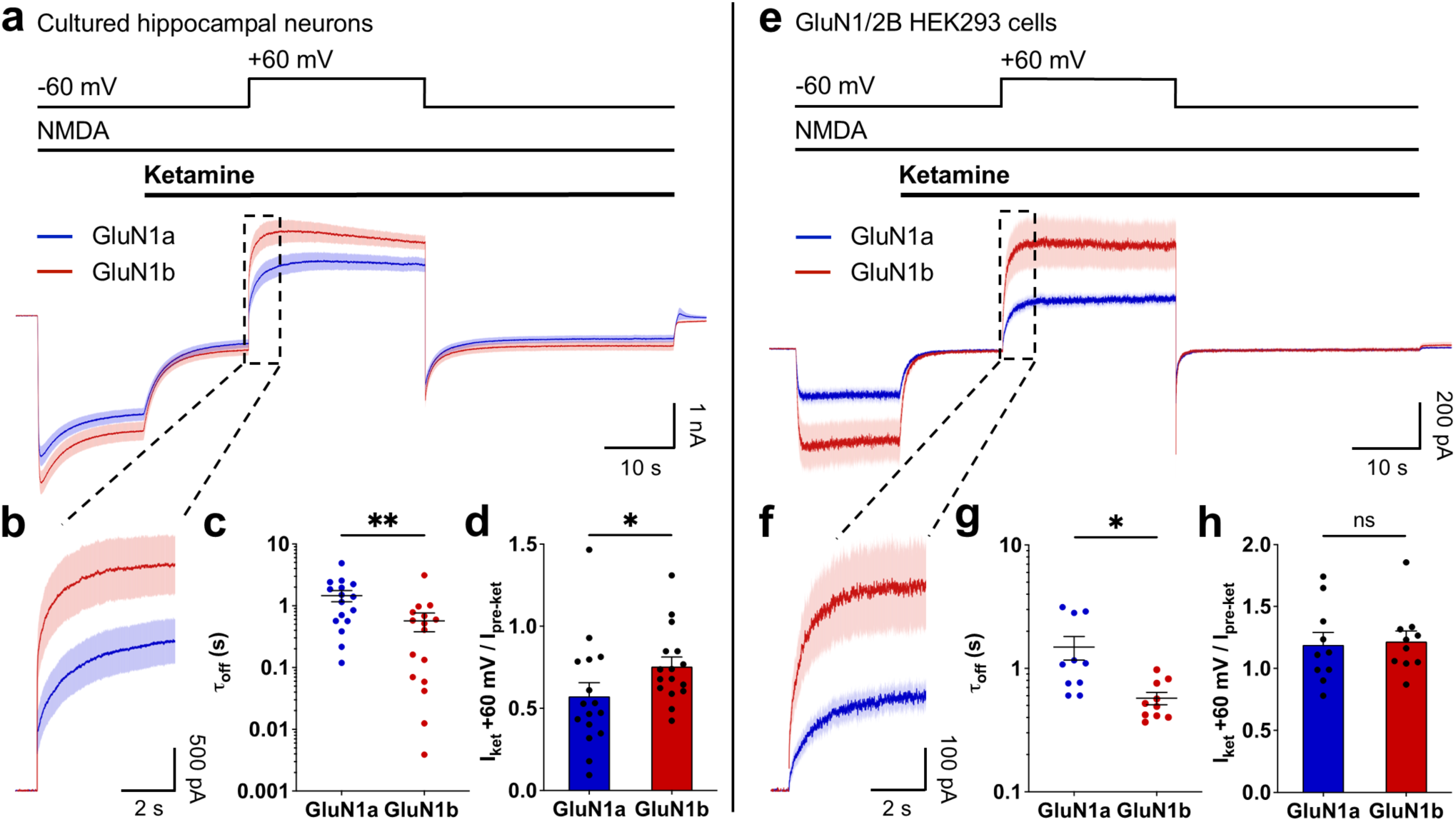
Voltage-dependent relief of ketamine inhibition is slower in GluN1a- than in GluN1b-containing NMDARs. NMDA-evoked currents with the membrane potential held at −60 mV in the absence of extracellular Mg^2+^. Ketamine (10 μM) was applied during NMDA administration. The membrane potential was depolarized to +60 mV for 25 s, to trigger ketamine unblocking, and repolarized to −60 mV. **a,** NMDA-evoked currents during the depolarization protocol in hippocampal neurons cultured from GluN1a (n = 16) and GluN1b (n = 16) mice. **b,** Currents from **(a)** during the initial 5 s upon depolarization. Currents prior to the depolarizing step were aligned for comparison in GluN1a and GluN1b neurons. **c,** Tau off of ketamine upon depolarization. **d,** Peak amplitude of NMDA-evoked currents at +60 mV during ketamine application relative to the current at −60 mV before ketamine application. **e,** NMDA-evoked currents during the depolarization protocol in HEK293 cells expressing GluN1a (n = 10) or GluN1b (n = 10) with GluN2B. **f,** Currents from **(e)** during the initial 5 s upon depolarization. Currents prior to the depolarizing step were aligned for comparison in GluN1a and GluN1b cells. **g,** Tau off of ketamine upon depolarization. **h,** Peak amplitude of NMDA-evoked currents at +60 mV during ketamine application relative to the current at −60 mV before ketamine application. Data are represented as mean ± SEM. *p < 0.05, **p < 0.01, ns p > 0.05, Student’s t-test or Mann-Whitney test, as appropriate.

To interrogate the effect of the N1 cassette on the voltage-dependent relief of ketamine while experimentally controlling the subunit composition, we used recombinant NMDARs expressed in HEK293 cells (GluN1a or GluN1b together with GluN2B) (Fig. 3E, F). In these recordings, the maximum current at +60 mV in GluN1a cells did not differ from that in GluN1b cells (relative to pre-ketamine current, p = 0.684; Fig. 3H). Similar to the cultured neurons, we found that the rate of unblock upon depolarization was 2.6-fold slower in GluN1a than in GluN1b cells (GluN1a τ = 1.49 ± 0.33 s, GluN1b τ = 0.575 ± 0.066 s, p = 0.013; Fig. 3G). Overall, we conclude that inclusion of the N1 cassette markedly accelerates the rate of voltage-dependent relief of ketamine blockade.

### Ketamine block is relieved during bursting activity from NMDARs containing the N1 cassette

To explore how the difference in the voltage-dependent ketamine effects imparted by the N1 cassette may affect NMDAR currents during TBS, we first mimicked TBS-patterned depolarization in recordings from hippocampal cultured neurons. During ketamine blockade of NMDA-evoked currents, we applied a TBS-like train comprised of 15 x 50 ms depolarizing steps from −60 mV to +60 mV at 5 Hz (Fig. 4A, B). During depolarization to +60 mV, we found in GluN1a neurons compared to in GluN1b neurons: significantly smaller current amplitudes (GluN1a = 0.458 ± 0.068, GluN1b = 0.681 ± 0.070 of pre-ketamine current, p = 0.0295; Fig. 4C, D) and less charge transfer (GluN1a = 0.398 ± 0.063, GluN1b = 0.585 ± 0.065 of pre-ketamine current, p = 0.0292; Fig. 4C, E). Furthermore, during repolarization to −60 mV, we observed substantial reblocking in GluN1b neurons (Fig. 4F-H), while reblocking was minimal in GluN1a neurons. Thus, during TBS-patterned depolarization, ketamine blockade is relieved upon depolarization and re-established upon repolarization in GluN1b-containing NMDARs, whereas ketamine blockade is largely retained in GluN1a-containing NMDARs.

**Figure 4.**
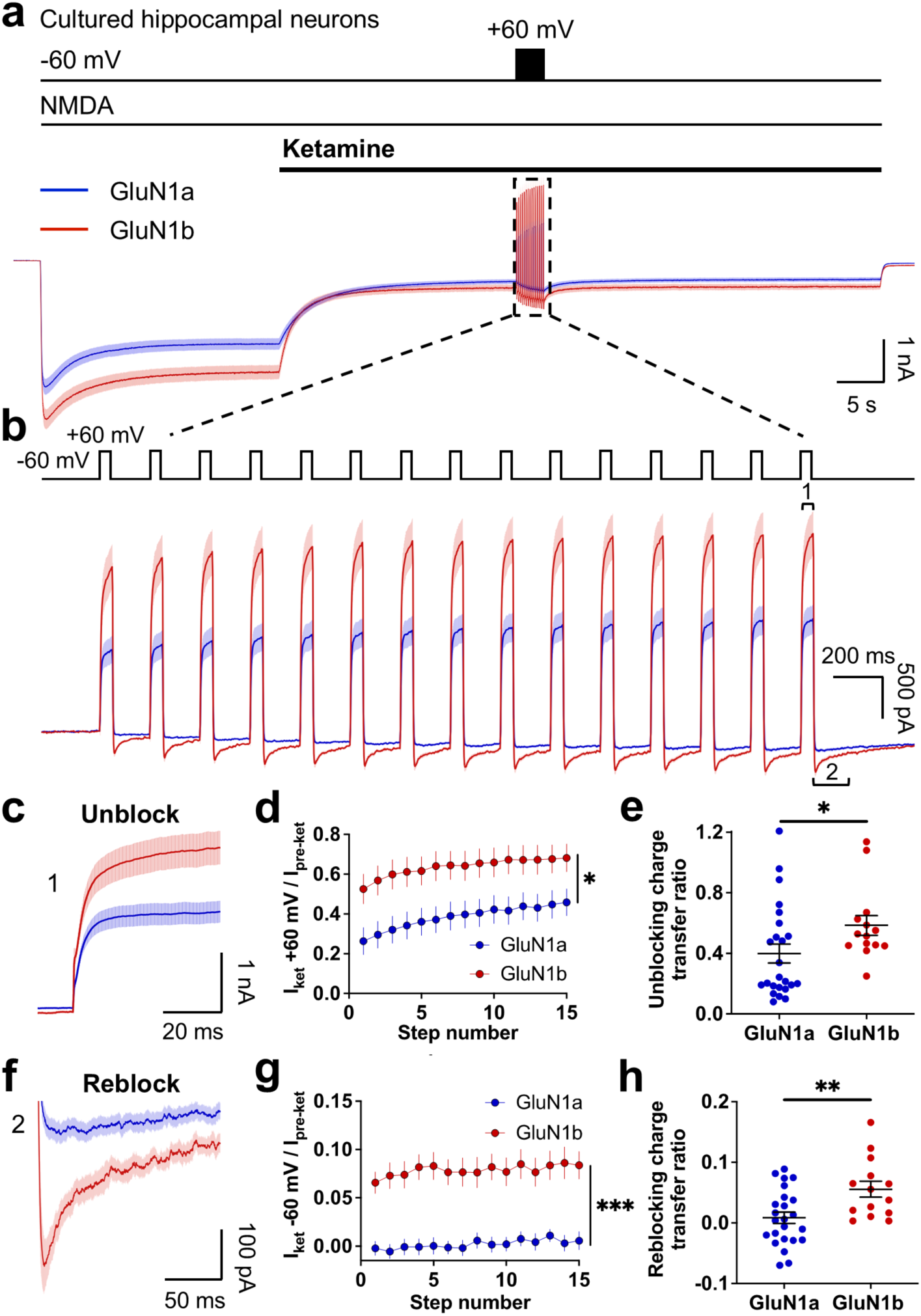
Relief of ketamine inhibition during theta-burst patterned stimulation is less in GluN1a- than in GluN1b-containing NMDARs. **a,** NMDA-evoked currents in hippocampal neurons cultured from GluN1a (n = 24) and GluN1b (n = 14) mice. The membrane potential was held at −60 mV. Ketamine (10 μM) was applied during NMDA administration, followed by a TBS-patterned protocol (15 x 50 ms depolarizing steps from −60 mV to +60 mV at 5 Hz). **b,** Currents from **(a)** during the TBS-patterned protocol. Currents prior to the first depolarizing step were aligned for comparison in GluN1a and GluN1b neurons. **c,** Currents from **(b)** during time point 1. Peak amplitude of NMDA-evoked currents at +60 mV **(d)** and charge transfer of the unblock during the last depolarizing step **(e)** relative to that for the current at −60 mV before ketamine application. **f,** Currents from **(b)** during time point 2. **g,** Peak to end amplitude of NMDA-evoked currents at −60 mV relative to the amplitude of the current before ketamine application. **h,** Charge transfer of the reblock during the last repolarizing step relative to that for the total current during the step. Data are represented as mean ± SEM. *p < 0.05, **p < 0.01, ***p < 0.001, two-way repeated measures ANOVA, Student’s t-test or Mann-Whitney test, as appropriate.

From these findings, we predicted that the synaptic responses during TBS, which contain a substantial NMDAR-dependent component^31,33^, are differentially affected by ketamine in GluN1a and GluN1b neurons. We made current-clamp recordings from CA1 pyramidal neurons in hippocampal slices and evoked postsynaptic potentials (PSPs) by stimulating the Schaffer collaterals. Slices were either pre-treated with ketamine for 1 h or untreated in controls. Ketamine treatment prevented LTP in GluN1a neurons but had no effect on LTP in GluN1b neurons (GluN1a = 105 ± 3.9%, GluN1b = 194 ± 13%, p < 0.0001; Fig. 5A, B), corroborating our findings from the extracellular field recordings. During TBS, each four-stimulus burst elicited a rapid onset depolarizing envelope and triggered spiking, with the membrane potential then decaying prior to the next burst (Extended Fig. 5A). The NMDAR-dependent component of the depolarizing envelope was identified by applying the NMDAR antagonist D-APV (50 uM), which reduced the peak amplitude and accelerated the decay phase for each burst (Extended Fig. 6). We found in GluN1a neurons treated with ketamine, the peak amplitude was smaller and the decay was faster for each burst compared to that in GluN1b neurons treated with ketamine (Fig. 5C). In contrast, the depolarizing envelopes were not different between GluN1a and GluN1b neurons in untreated controls (Fig. 5D). We determined that ketamine significantly reduced the NMDAR-dependent component in GluN1a neurons but not in GluN1b neurons (Fig. 5E). Spike number was not different between GluN1a and GluN1b neurons treated with ketamine (Extended Fig. 5B). Taking our findings together, we conclude that during TBS, inclusion of the N1 cassette enables NMDARs to escape blockade by ketamine (Fig. 2A). By contrast, ketamine blockade persists during TBS in NMDARs lacking the N1 cassette.

**Figure 5.**
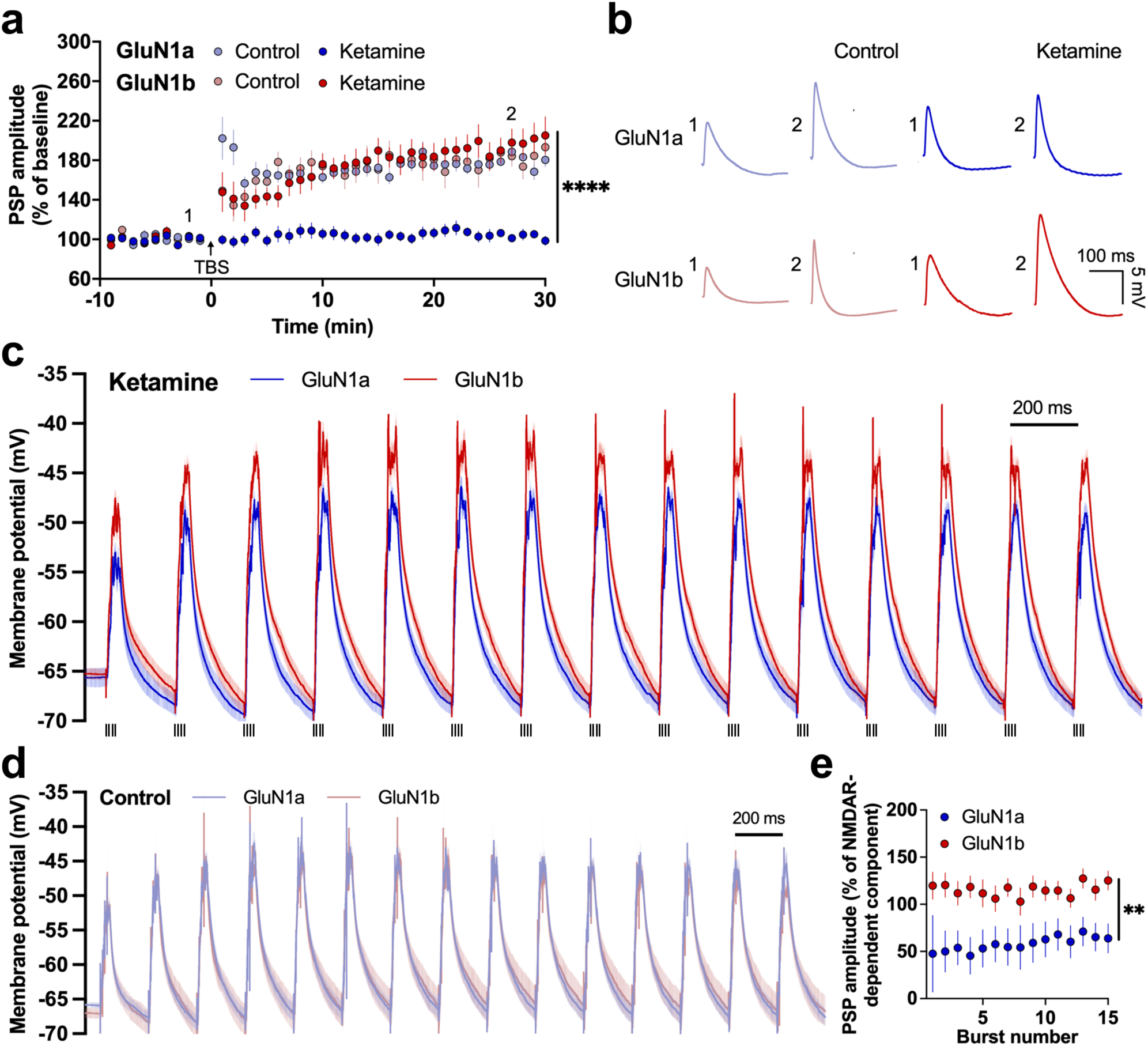
Ketamine blockade is relieved during bursting activity in GluN1b- but not in GluN1a-containing NMDARs. **a,** Time plot of PSP amplitudes from CA1 pyramidal neurons in hippocampal slices from GluN1a (control n = 8, ketamine n = 7) and GluN1b (control n = 5, ketamine n = 8) mice. Slices were pre-treated with ketamine (10 µM) for 1 h or untreated in controls. TBS was delivered at time 0. PSP amplitudes are normalized to the baseline. **b,** Representative PSP traces at time points 1 and 2. Data are represented as mean ± SEM. ****p < 0.0001, two-way ANOVA with Tukey’s post hoc test. PSPs during TBS in GluN1a (n = 8) and GluN1b (n = 9) neurons from ketamine-treated slices **(c)** and in GluN1a (n = 9) and GluN1b (n = 6) neurons from control slices **(d)**. Action potentials were truncated to examine the underlying PSP amplitudes. **e,** PSP amplitudes in ketamine-treated relative to control neurons as a percentage of the NMDAR-dependent component identified by the effect of D-APV across bursts (Extended data Fig. 6). Data are represented as mean ± SEM. **p < 0.01, two-way repeated measures ANOVA.

### Antidepressant effect of ketamine is prevented by the presence of the N1 cassette

Based on the differential effects of ketamine on GluN1a-versus GluN1b-containing NMDARs, we investigated the possibility that the presence of the N1 cassette has consequences for the antidepressant activity of ketamine. We conducted the tail suspension test (TST), in which a reduction in immobility time is a highly validated predictive measure of the antidepressant activity of drugs, including ketamine^34^. We administered ketamine (10 mg/kg) or saline vehicle control via i.p. injection to GluN1a, GluN1b, and WT mice and performed the TST 1 h later (Fig. 6A). WT mice treated with ketamine exhibited significantly less immobility than their saline-treated controls (saline = 271 ± 6.3 s, ketamine = 237 ± 6.8 s, p = 0.0003; Fig. 6A), as expected. We found that GluN1a mice treated with ketamine were also significantly less immobile than their respective saline-treated controls (saline = 267 ± 3.7 s, ketamine = 230 ± 9.1 s, p = 0.0005; Fig. 6A). In contrast, immobility levels in GluN1b mice treated with ketamine were not different from those treated with saline (saline = 270 ± 6.6 s, ketamine = 270 ± 5.7 s, p = 0.960; Fig. 6A). Furthermore, immobility levels in ketamine-treated WT or GluN1a mice were also significantly less than that in ketamine-treated GluN1b mice (p = 0.002, p = 0.0005, respectively; Fig. 6A). We therefore conclude that inclusion of the N1 cassette prevents the antidepressant effect of systemic ketamine treatment.

**Figure 6.**
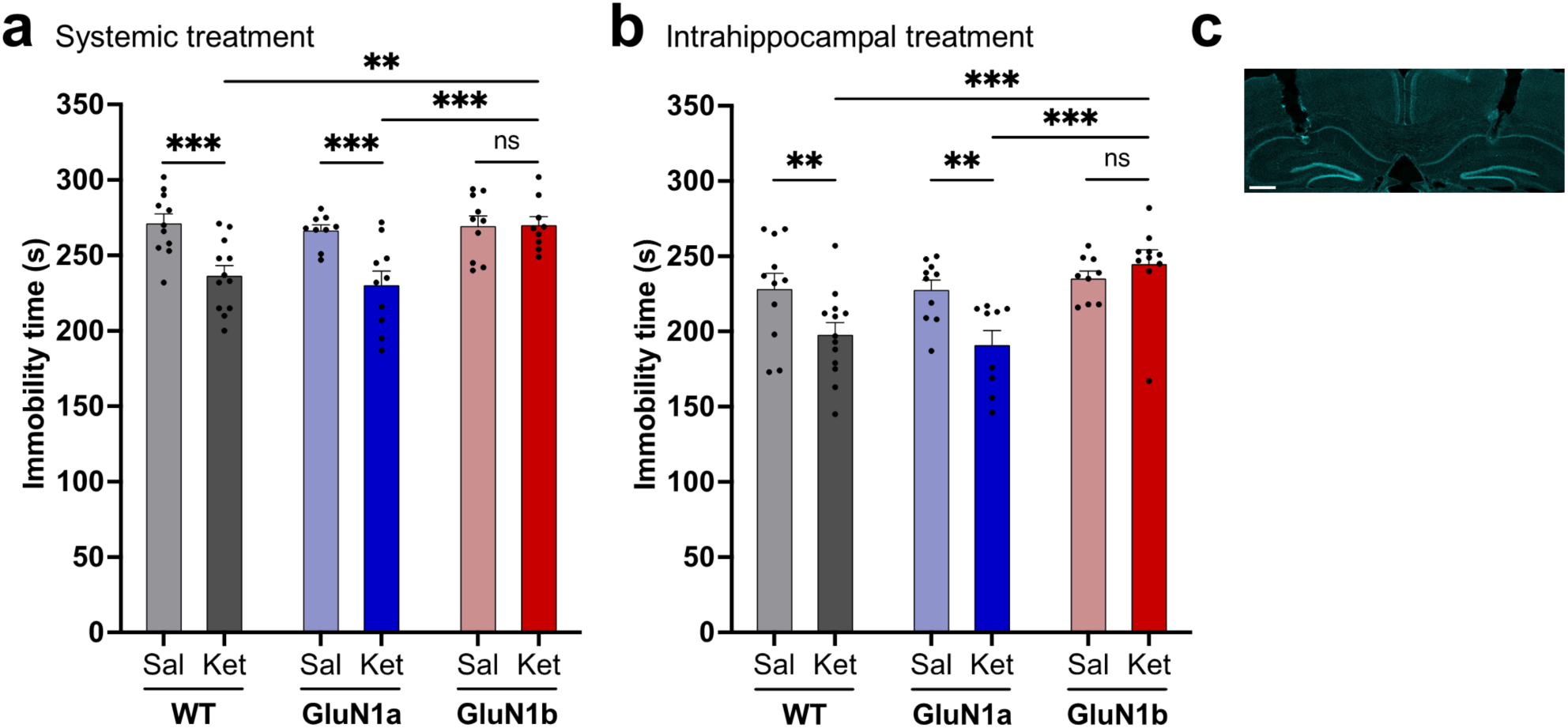
Ketamine induces an antidepressant effect in GluN1a mice but not in GluN1b mice. **a,** Ketamine (ket) (10 mg/kg) or saline (sal) was administered by i.p. injection to WT (sal n = 11, ket n = 12), GluN1a (sal n = 9, ket n = 10), and GluN1b (sal n = 10, ket n = 9) mice. Immobility levels were measured 1 h later in the TST. **b,** Bilateral hippocampal infusions of ketamine (3 μg) or saline were performed in WT (sal n = 11, ket n = 13), GluN1a (sal n = 10, ket n = 9), and GluN1b (sal n = 9, ket n = 10) mice. Immobility levels were assessed 1 h later in the TST. **c,** Bilateral cannula sites in the hippocampus. DAPI in cyan. Scale bar = 500 μm. Data are represented as mean ± SEM. **p < 0.01, ***p < 0.001, ns p > 0.05, two-way ANOVA with Tukey’s post hoc test.

The hippocampus has been strongly implicated in the antidepressant effects of ketamine^10,16,35–38^. We found that bilateral infusion of ketamine (3 μg in 1 μL) into the hippocampus (Fig. 6C) significantly reduced immobility 1 h later in WT mice (saline = 228 ± 10 s, ketamine = 198 ± 8.0 s, p = 0.0091; Fig. 6B), demonstrating that infusion of ketamine into the hippocampus was sufficient to produce its antidepressant effect. GluN1a mice infused with ketamine were also significantly less immobile than their saline-treated controls (saline = 228 ± 6.6 s, ketamine = 191 ± 10 s, p = 0.0054; Fig. 6B). However, GluN1b mice infused with ketamine were not different in immobility than their respective controls (saline = 235 ± 5.0 s, ketamine = 245 ± 9.4 s, p = 0.447; Fig. 6B). Immobility levels were also significantly less in WT or GluN1a mice than that in GluN1b mice infused with ketamine (p = 0.0004, p = 0.0002, respectively; Fig. 6B). Thus, inclusion of the N1 cassette also prevents the antidepressant effect induced by intrahippocampally administered ketamine.

## Discussion

Here we demonstrate that the exclusion or inclusion of the N1 cassette, in a region of the receptor distant from the channel pore, has striking consequences for the dynamics of NMDAR blockade and antidepressant action of ketamine. An important consideration in interpreting our findings is that the replacement strategy we used did not cause alterations in the expression of *Grin1* or the amplitude of synaptic NMDAR currents^18^. Using this strategy, we avoid a major confound in previous studies that investigated the role of NMDARs in the antidepressant effect of ketamine, in which genomic deletion of NMDAR subunits reduced the total amount of NMDARs and produced marked alterations in basal synaptic function, neuronal excitability, and behaviour^11,35^. In the present study, eliminating GluN1a-containing NMDARs, by replacing them with GluN1b-containing NMDARs in GluN1b mice, abolished the antidepressant effect of ketamine. Conversely, the antidepressant effect was intact in GluN1a mice, which only expressed GluN1a-containing NMDARs. We therefore conclude that GluN1a-containing NMDARs are both necessary and sufficient for the antidepressant activity of ketamine.

We found that the steady-state level of ketamine blockade in synaptic GluN1a-containing NMDARs was around 20% greater than that in GluN1b-containing NMDARs holding at a negative membrane potential, which might contribute in part to the requirement of GluN1a-containing NMDARs for the antidepressant effect. Moreover, the rate at which voltage-dependent relief of ketamine blockade develops is several-fold slower for GluN1a- than for GluN1b-containing receptors. Consequently, during brief periods of depolarization as observed in TBS, a physiological activity pattern of hippocampal pyramidal neurons in behaving animals^26,27,39^, GluN1b-containing NMDARs escape ketamine blockade whereas GluN1a-containing NMDARs, unable to escape, remain blocked. Thus, we conclude that the antidepressant action of ketamine is not mediated by indiscriminate blockade of all NMDARs. Instead, the antidepressant action of ketamine is mediated selectively by GluN1a-containing NMDARs, in which the rate of voltage-dependent unblock is sufficiently slow to satisfy the dynamics required to maintain ketamine blockade during neuronal firing activity (Fig. 7).

**Figure 7.**
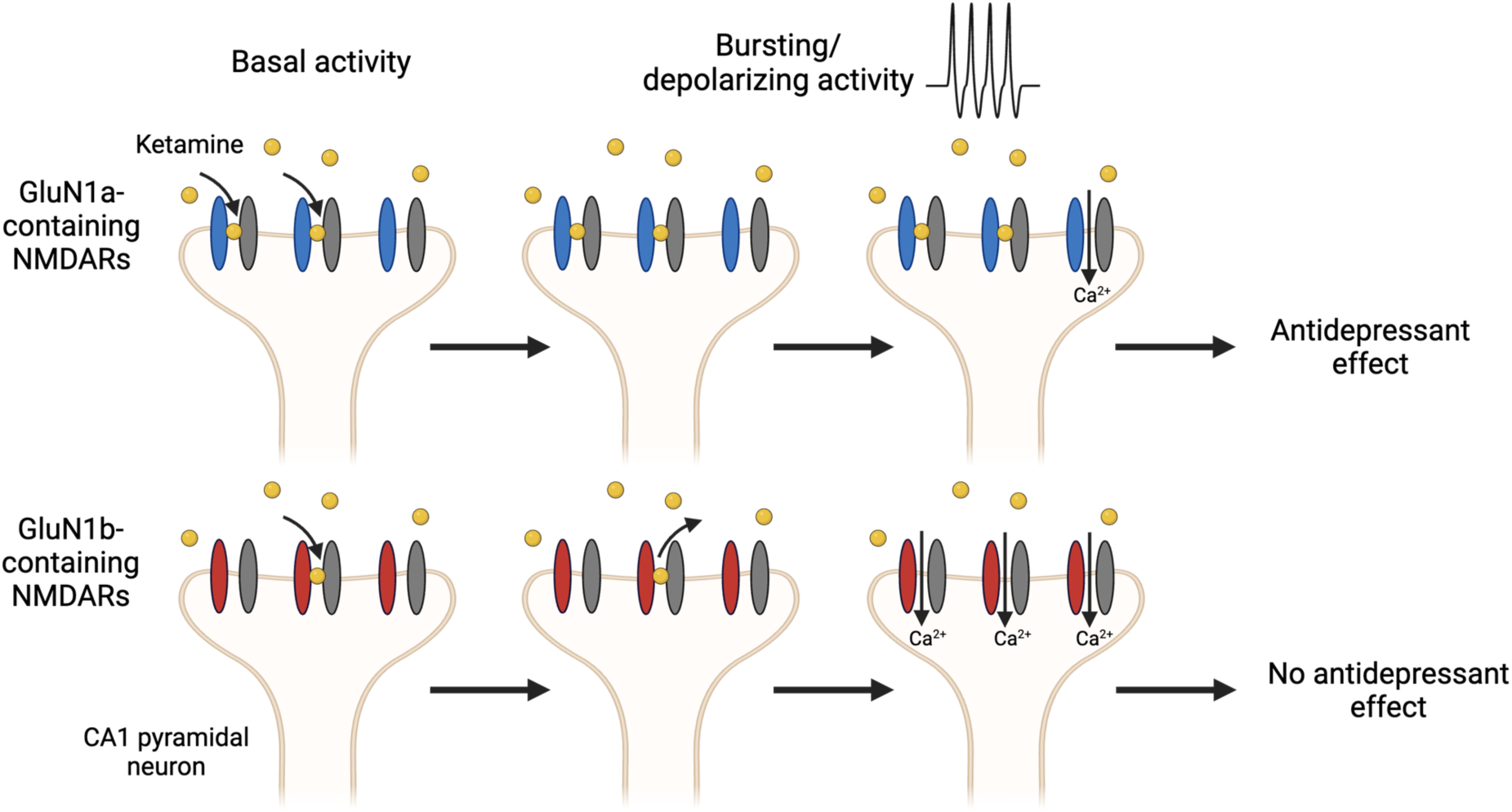
Ketamine blockade of GluN1a-containing NMDARs mediates its antidepressant activity. Ketamine preferentially inhibits synaptic GluN1a-containing NMDARs over those containing GluN1b during basal activity. Upon high frequency depolarizing activity, ketamine blockade is relieved from GluN1b-containing receptors, while the blockade persists in GluN1a-containing receptors, limiting current flow through these receptors. This persistent blockade of GluN1a-containing receptors drives the antidepressant effect of ketamine.

The N1 cassette is located in the extracellular N-terminal domain^22^, too far away from the channel pore to directly affect the site where ketamine binds within the pore^40,41^. Thus, the N1 cassette must be acting at a distance to control the rate at which depolarization drives ketamine from the channel. The N1 cassette is known to interact with residues at the interface of ligand binding domains of GluN1 and GluN2^22^, which affect allosteric modulation and deactivation of the receptor^42–44^. Therefore, it is conceivable that through these interactions, the N1 cassette accelerates the voltage-dependent relief of ketamine blockade.

Our observations that direct intrahippocampal administration of ketamine is sufficient to produce antidepressant effects in both GluN1a and WT mice but has no effect in GluN1b mice, comparable to the effects of systemic administration, is in line with the higher relative expression of GluN1a- over GluN1b- containing NMDARs in the hippocampus of rodents^19,45,46^. In the human brain, global expression of GluN1a and GluN1b transcripts are typically around 50% each^20,21^, but there is considerable regional variation. The hippocampus predominantly expresses GluN1a over GluN1b compared to other brain regions^20^. Collectively, these observations provide a basis for hypothesizing that the antidepressant action of ketamine may be mediated, at least in part, through blockade of GluN1a-containing NMDARs in the hippocampus. As GluN1a-containing NMDARs are also expressed in other brain regions, our study opens up the possibility that blockade of GluN1a-containing NMDARs in regions beyond the hippocampus might contribute to the antidepressant effect of ketamine.

NMDARs are long-standing therapeutic targets for numerous central nervous system disorders^47,48^. Development of NMDAR-targeting therapeutics has been overwhelmingly focused on selectivity for GluN2 subtypes^48,49^. However, our findings that the N1 cassette unexpectedly gates the actions of ketamine bring to light the importance of considering GluN1 and its alternative splicing in the actions of current NMDAR-targeting pharmacological treatments and in the future development of novel therapeutics.

## Acknowledgements

We sincerely thank Janice Hicks and Yongqian Wang for the technical support, as well as Dr. Graham Pitcher and Dr. Adam Fekete for the scientific discussions. This study was supported by funding from the Canadian Institutes of Health Research (CIHR) (FDN-154336 and PJT-191976) and the Krembil Foundation to M.W.S.. A.T.H. was supported by scholarships from CIHR and the Restracomp Scholarship from The Hospital for Sick Children.

## Author contributions

Conceptualization, A.T.H. and M.W.S.; Methodology, A.T.H. and M.W.S.; Formal Analysis, A.T.H., W.Z., and H.L.; Investigation, A.T.H., W.Z., H.L., Y.T., D.L., and D.K.; Writing – original draft, A.T.H. and M.W.S.; Writing – review and editing, A.T.H., W.Z., H.L., D.K., Z.J., L.Y.W., and M.W.S.; Visualization, A.T.H. and W.Z.; Supervision, M.W.S.; Funding Acquisition, M.W.S.

## Methods

### Animals

GluN1a and GluN1b mice were generated as previously described^18^. Mice were group housed under a 14:10 h light-dark cycle with access to food and water *ad libitum*. Experimental use of animals was in accordance with policies from the Hospital for Sick Children Animal Care Committee and the Canadian Council on Animal Care.

### Drugs

Ketamine (Ketamine hydrochloride, Toronto Research Chemicals) was bath applied in aCSF in electrophysiological experiments. Ketamine was dissolved in 0.9% saline and administered by i.p. injection or intrahippocampal infusion before behavior experiments.

### Brain slice preparation

Slice electrophysiology was performed in 4-5-week-old male and female mice, and the data are reported in aggregate. Mice were anesthetized with an intraperitoneal injection of 20% urethane. The brain was removed in ice-cold aCSF containing 124 mM NaCl, 3 mM KCl, 1.25 mM NaH_2_PO_4_, 2 mM MgCl_2_, 2 mM CaCl_2_, 23 mM NaHCO_3_, and 11 mM D-glucose saturated with 95% O_2_/5% CO_2_ (pH 7.40, 305 mOsm). Sagittal slices (300-μm thick) were obtained in ice-cold oxygenated aCSF using a vibratome. Slices were incubated in aCSF saturated with 95% O_2_/5% CO_2_ at 32°C for 40 min and placed at room temperature (22-24°C) for at least 1 h before recording. Slices were transferred to a submerged recording chamber perfused with aCSF saturated with 95% O_2_/5% CO_2_ at a flow rate of 2 mL/min at 27-28°C.

### Field recordings in hippocampal slices

Schaffer collateral afferents were stimulated with a bipolar tungsten electrode. Testing stimuli were 0.1 ms in duration and delivered at a frequency of 0.033 Hz to evoke a half-maximal fEPSP. Field excitatory postsynaptic potentials (fEPSPs) were recorded using an aCSF-filled glass micropipette. Ketamine (10 µM) was bath applied to slices for 1 h. LTP was induced using theta-burst stimulation (TBS) consisting of 15 bursts at 5 Hz, with each burst comprised of 4 pulses at 100 Hz. fEPSP slope was calculated from 10-60% of the peak amplitude. Signals were amplified with a MultiClamp 700B amplifier and digitized using a Digidata 1440A acquisition system sampled at 10 kHz. Data was analyzed using Clampfit 10.7. The experimenter was blinded to the genotype of the mice.

### Whole-cell recordings in hippocampal slices

Schaffer collateral afferents were stimulated with a bipolar tungsten electrode. Testing stimuli were 0.1 ms in duration and delivered at a frequency of 0.1 Hz. CA1 pyramidal neurons were visualized on a microscope (Zeiss Axioskop 2FS microscope). Pipettes had a typical resistance of 5–8 MΩ. In voltage-clamp recordings, NMDAR excitatory postsynaptic currents (EPSCs) were pharmacologically isolated with NBQX (10 μM) to block AMPA receptors and bicuculline (10 μM) to block GABA_A_ receptors. NMDAR EPSCs were recorded with an intracellular solution containing 117 mM Cs-gluconate, 10 mM CsCl_2_, 10 mM BAPTA, 1 mM CaCl_2_, 10 mM HEPES, 2 mM Mg-ATP, 10 mM QX-314, and 0.3 mM GTP (pH 7.25, 290 mOsm). After whole-cell configuration, the membrane potential was held at −40 mV in Mg^2+^-containing aCSF. Ketamine (10 µM) was bath applied to slices for 30 min. NMDAR EPSC signals were filtered at 1 kHz and peak amplitudes were measured. In current-clamp recordings, postsynaptic potentials (PSPs) were recorded with an intracellular solution containing 130 mM K-gluconate, 10 mM KCl, 0.2 mM EGTA, 10 mM HEPES, 4 mM Mg-ATP, 10 mM Na phosphocreatine, and 0.3 mM GTP (pH 7.25, 290 mOsm). Slices were pre-treated with ketamine (10 µM) for 1 h before recording. TBS was applied to induce LTP (15-16 bursts at 5 Hz, with each burst comprised of 4 pulses at 100 Hz). D-APV (50 µM) treatment for 20 min was used to identify the NMDAR-dependent component of the PSP during TBS. Peak amplitudes of single PSPs were measured in LTP recordings. Peak amplitudes underlying the action potentials were measured during TBS. Spike number was also calculated per burst. Signals were amplified with a MultiClamp 700B amplifier and digitized using a Digidata 1440A acquisition system sampled at 10 kHz. Data was analyzed using Clampfit 10.7.

### Hippocampal neuronal culture preparation

Hippocampal neuronal cultures were prepared from timed pregnant mice. The brain was removed from E15 fetuses in ice-cold Hank’s solution. The hippocampus was dissected from each hemisphere and pooled for each embryo. Each embryo’s pooled hippocampi were mechanically dissociated through a 1000 μL pipette tip. The cells were plated onto poly-D-lysine-coated glass coverslips. The culture media was Neurobasal medium supplemented with fetal bovine serum, L-glutamine, and B-27 supplement. Electrophysiological recordings were performed after 13-19 d in culture at room temperature (22°C).

### HEK293 cell preparation

Human embryonic kidney (HEK293) cells were cultured in Dulbecco’s Modified Eagles Medium (DMEM) supplemented with 10% fetal bovine serum (FBS) and 1% penicillin-streptomycin at 37°C in a 5% CO_2_ incubator. HEK293 cells were seeded on 35×10 mm dishes at 4 x 10^5^ cells per dish 24 h before transfection. FuGene HD (Promega BioSciences) was used for all transfections. Transfections contained expression plasmids encoding GluN1-1a or GluN1-1b, GluN2B, PSD-95, and a fluorescent protein (green fluorescent protein or tdTomato) at a ratio of 1:4:0.5:0.5. After transfection, cells were maintained in DMEM supplemented with 10% FBS and 500 μM D-APV for 48 h before electrophysiological recordings were performed at room temperature (22°C).

### Whole-cell recordings in hippocampal cultured neurons and HEK293 cells

Whole-cell voltage-clamp recordings were made in an extracellular recording solution composed of 140 mM NaCl, 5.4 mM KCl, 15 mM HEPES, 25 mM glucose, 1.3 mM CaCl2, 0.0005 mM tetrodotoxin, and 0.001 mM glycine (pH 7.35). Tetrodotoxin was not included in HEK293 cell recordings. Pipettes had a typical resistance of 3-6 MΩ and were filled with an intracellular solution containing 137 mM CsF, 1.5 mM CsCl, 10 mM BAPTA, 10 mM HEPES, and 4 mM Mg-ATP (pH 7.20). After whole-cell configuration, the membrane potential was held at −40 mV in the presence of 1 mM extracellular Mg^2+^ or −60 mV in the absence of extracellular Mg^2+^. NMDAR currents were evoked by NMDA (50 µM) application using a fast-step perfusion system (SF-77B, Warner Instruments, USA). Ketamine (10 μM) was locally applied using this fast-step perfusion system during NMDA administration. In unblocking experiments, the membrane potential was stepped to +60 mV. The NMDAR component was obtained by subtracting the total transmembrane currents recorded by that evoked by the voltage step alone. Current amplitudes were measured and time constants were calculated from the time required for currents to reach 63.2% of their maximum value. In ketamine unblocking experiments in the presence of extracellular Mg^2+^, the starting point was 10 ms after the voltage step to minimize the contribution of the fast Mg^2+^ unblock^50^. A TBS-like protocol was applied consisting of 15 x 50 ms depolarizing steps from −60 mV to +60 mV at 5 Hz. The unblock was measured using the peak amplitude of NMDAR current at each step to +60 mV normalized to the steady-state current amplitude before ketamine application, and the unblocking charge transfer during the last 50 ms depolarization step was normalized to the charge transfer during 50 ms of the pre-ketamine current. The reblock was measured from the peak to end current amplitude during each step to −60 mV normalized to the steady-state current amplitude before ketamine application, and the reblock charge transfer during the last 150 ms repolarization step was normalized to the total charge transfer during the 150 ms repolarizing step. Signals were amplified using an Axopatch 1D amplifier and digitized using an Axon Digidata 1440A acquisition system. Signals were sampled at 5 kHz and filtered at 2 kHz. Data was analyzed using Clampfit 10.7. The experimenter was blinded to the genotype of the cells.

### Hippocampal cannula implantation and infusion

Mice were anesthetized with 3% isoflurane and placed in a stereotaxic frame (RWD instruments). Bilateral guide cannulas (RWD instruments) were implanted into the hippocampus (−2.2 mm AP, ±1.5 mm ML, −1.5 mm DV) of male mice. Dummy cannulas were inserted into the guide cannulas to prevent blockage. After a recovery period of 8 d, mice were anesthetized with isoflurane and internal cannulas were first inserted and removed to clear the infusion pathway. Ketamine dissolved in 0.9% saline or saline alone was infused using a microinjector pump (Harvard Apparatus). Ketamine (3 μg) or saline in 1 μL was infused at 0.5 μL/min 1 h before behavior experiments. To check the cannula implantation site, brain samples were post-fixed in formalin and 30% sucrose following perfusion. Subsequently, brains were cryosectioned into 50 µm slices followed by 10 min DAPI staining (1/1500). Images were acquired with a Quorum spinning disk confocal microscope using Volocity (Ver 6.5.1) software.

### Tail suspension test

Behavior experiments were performed in 10-12-week-old male mice^11,36,38^. Mice were handled once per day for 5 consecutive days prior to testing day. Mice were suspended by the tail from a bar in a tail suspension apparatus (55 cm height x 15 cm width x 11.5 cm depth) for 6 min. Their movement was recorded, and immobility time was quantified. The experimenter was blinded to the genotype and treatment of the animals.

### Quantification and statistical analysis

Graphs were made using GraphPad Prism 10 or R. Statistical analyses were performed using GraphPad Prism 10. Ns for each experiment are detailed in the figure legends. Normality of the data was evaluated using the D’Agostino and Pearson test and the appropriate parametric or non-parametric test was used i.e. Student’s t-test or Mann-Whitney test, respectively, as detailed in the corresponding figure legends.

All statistical tests were two-tailed, and statistical significance was considered as p < 0.05. Repeated-measures two-way ANOVA or two-way ANOVA with Tukey’s multiple comparisons test were used as appropriate, as indicated in the corresponding figure legends.

---

**Extended data Figure 1 related to Figure 1.**
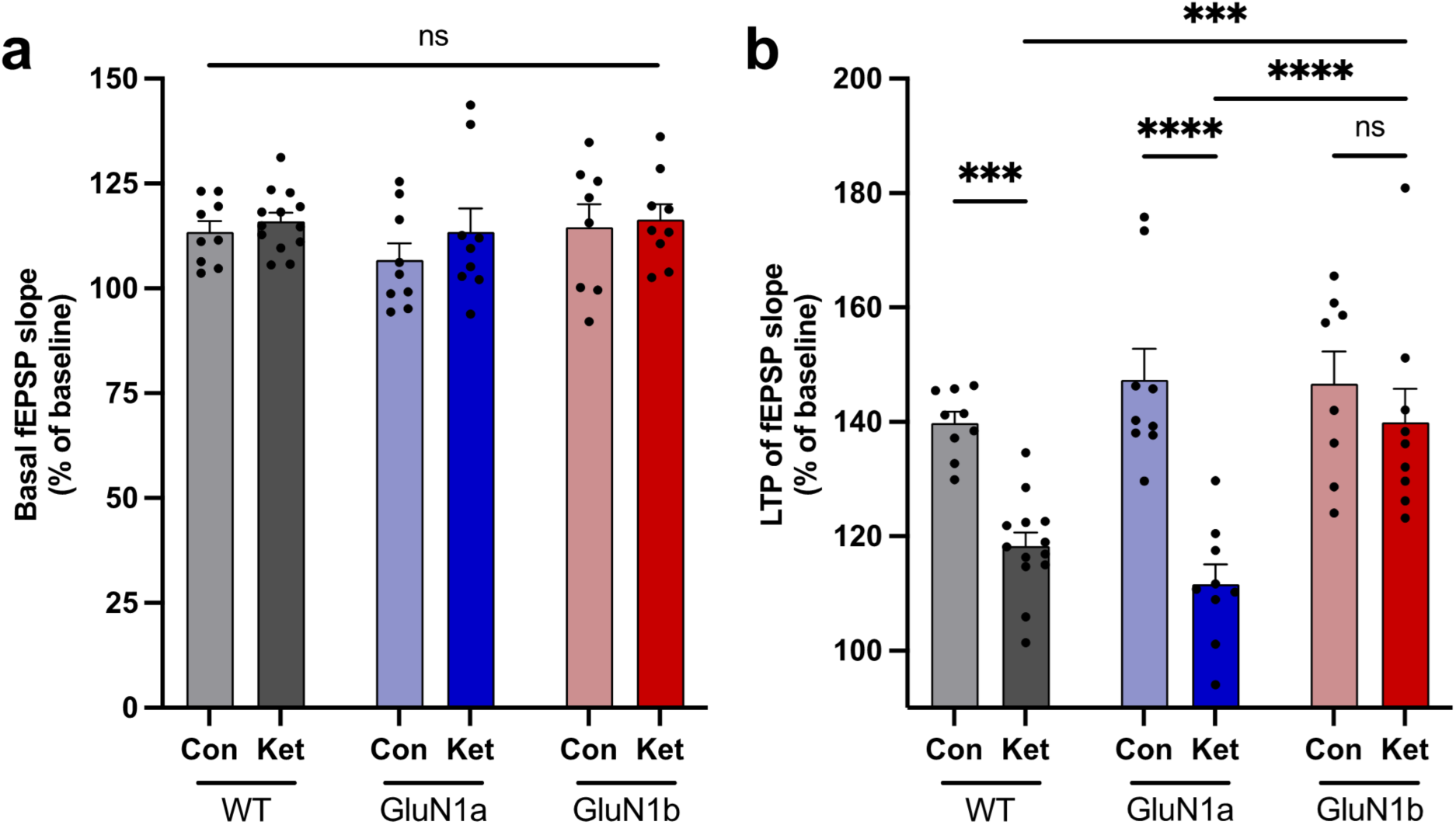
Ketamine inhibits LTP at Schaffer collateral-CA1 synapses in GluN1a but not in GluN1b hippocampal slices without affecting basal fEPSPs. **a,** fEPSP slopes of the last 10 min of 1 h ketamine (ket) (10 µM) treatment or no treatment control (con) normalized to baseline in WT (con n = 9, ket n = 13), GluN1a (con n = 9, ket n = 9), and GluN1b (con n = 8, ket n = 9) mice. **b,** LTP magnitude of the last 10 min of LTP normalized to the 10 min before TBS. Data are represented as mean ± SEM. ***p < 0.001, ****p < 0.0001, ns p > 0.05, two-way ANOVA with Tukey’s post hoc test.

**Extended data Figure 2 related to Figure 2.**
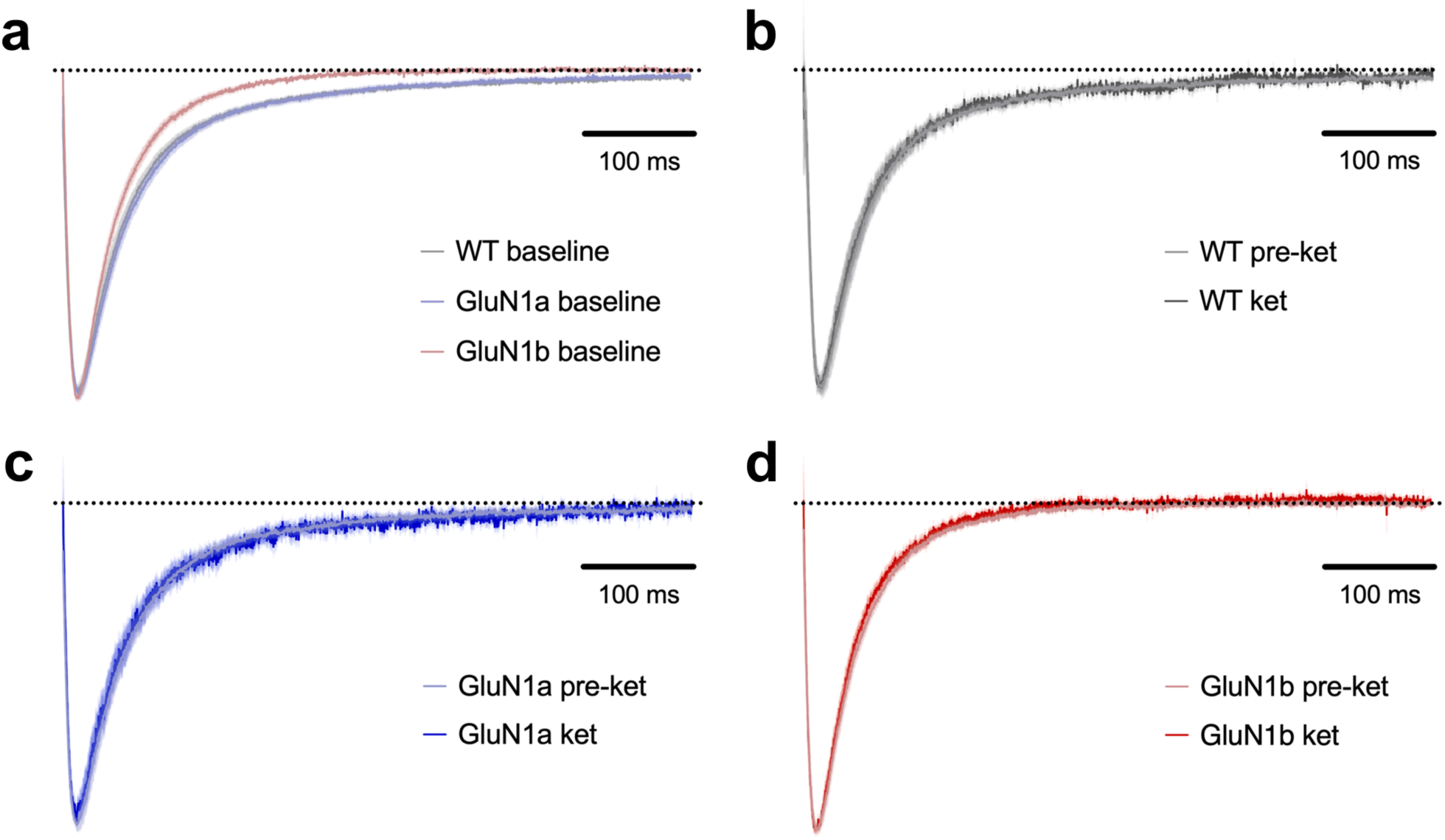
NMDAR EPSC rise and decay are unaffected by ketamine. **a,** NMDAR EPSCs from CA1 pyramidal neurons holding at −40 mV in hippocampal slices from WT (n = 15), GluN1a (n = 16), and GluN1b (n = 15) mice at baseline combined from untreated control and ketamine-treated slices. Note that NMDAR EPSC decay is slower in WT and GluN1a compared to in GluN1b mice, as previously reported^31^. NMDAR EPSCs following ketamine treatment (10 μM) for 30 min compared to baseline from **(b)** WT (n = 8), **(c)** GluN1a (n = 8), and **(d)** GluN1b (n = 8) mice. Data are scaled to peak to compare the rise and decay and represented as mean ± SEM.

**Extended data Figure 3 related to Figure 3.**
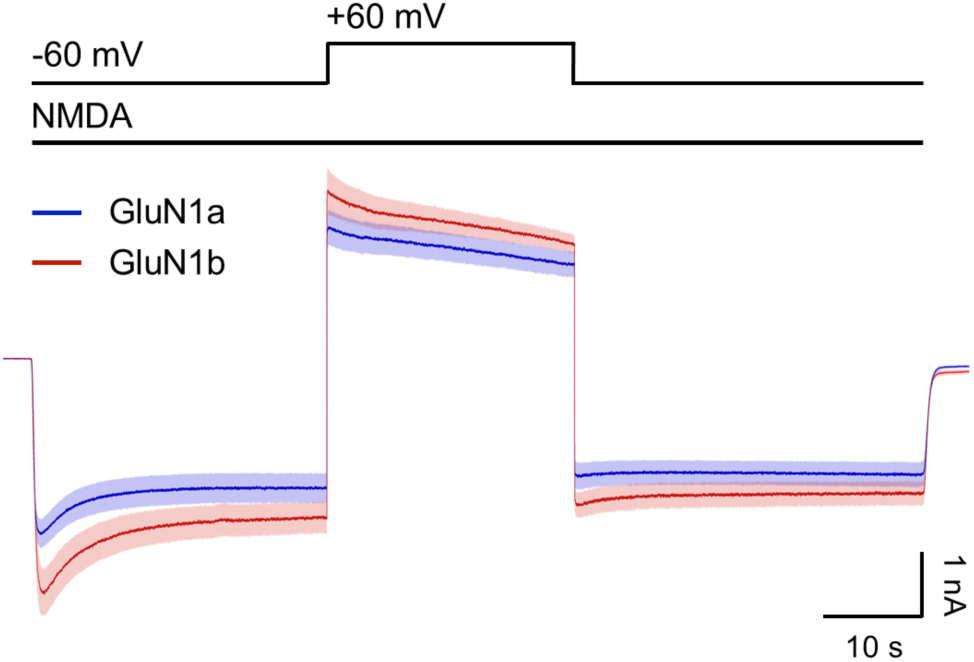
NMDAR currents are similar during depolarization in GluN1a and GluN1b neurons in the absence of ketamine. NMDA-evoked currents in hippocampal neurons cultured from GluN1a (n = 10) and GluN1b (n = 12) mice. The membrane potential was held at −60 mV in the absence of extracellular Mg^2+^. The membrane potential was depolarized to +60 mV for 25 s and repolarized to −60 mV. Data are represented as mean ± SEM.

**Extended data Figure 4 related to Figure 3.**
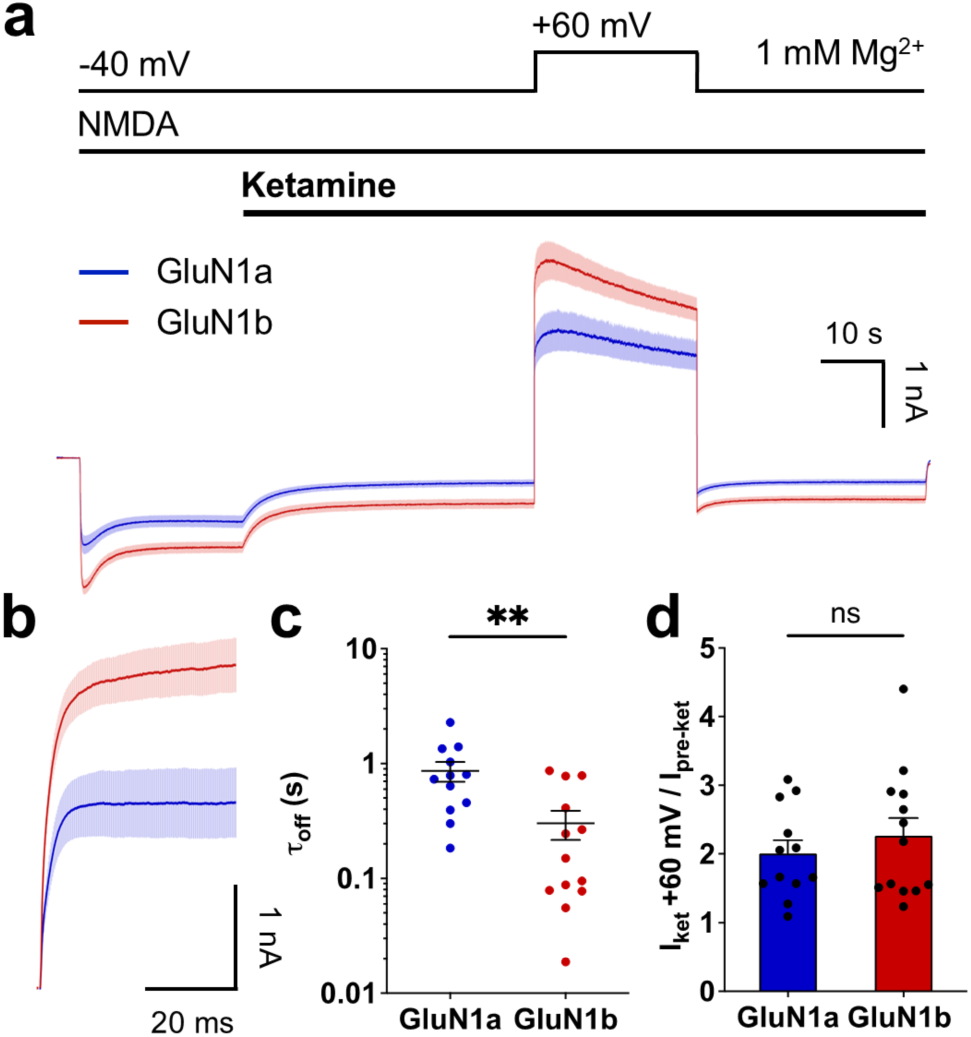
Voltage-dependent relief of ketamine inhibition is slower in GluN1a- than in GluN1b-containing NMDARs in the presence of extracellular Mg^2+^. **a,** NMDA-evoked currents in hippocampal neurons cultured from GluN1a (n = 12) and GluN1b (n = 13) mice. The membrane potential was held at −40 mV in the presence of 1 mM extracellular Mg^2+^. Ketamine (10 μM) was applied during NMDA administration. The membrane potential was depolarized to +60 mV for 25 s and repolarized to −40 mV. **b,** Currents from **(a)** during the initial 50 ms upon depolarization. Currents prior to the depolarizing step were aligned for comparison in GluN1a and GluN1b neurons. **c,** Tau off of ketamine upon depolarization. **d,** Peak amplitude of NMDA-evoked currents at +60 mV during ketamine application relative to the current at −40 mV before ketamine application. Data are represented as mean ± SEM. **p < 0.01, ns p > 0.05, Student’s t-test or Mann-Whitney test, as appropriate.

**Extended data Figure 5 related to Figure 5.**
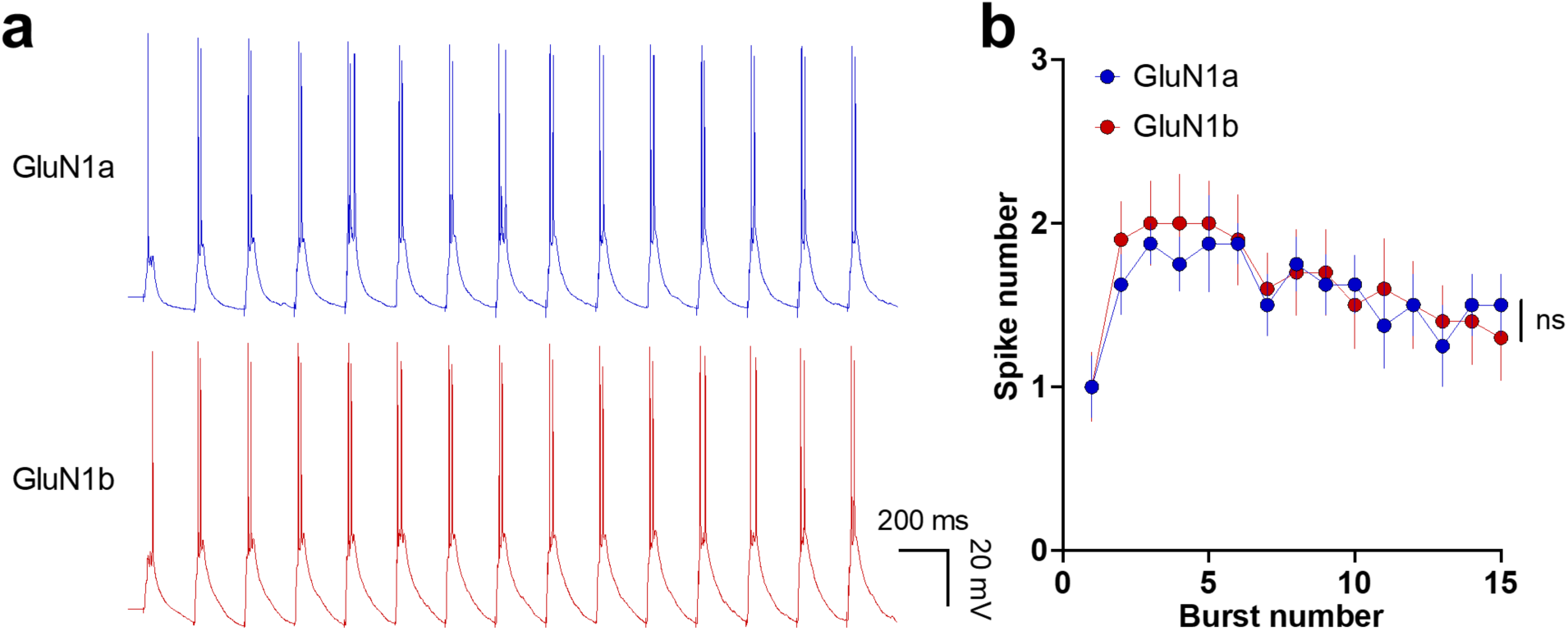
Spike number is not different between GluN1a and GluN1b neurons during bursting activity. **a,** Representative PSP traces during TBS from CA1 pyramidal neurons in hippocampal slices pre-treated with ketamine (10 µM) for 1 h from GluN1a (n = 8) and GluN1b (n = 9) mice. **b,** Spike number across bursts. Data are represented as mean ± SEM. ns p > 0.05, two-way repeated measures ANOVA.

**Extended data Figure 6 related to Figure 5.**
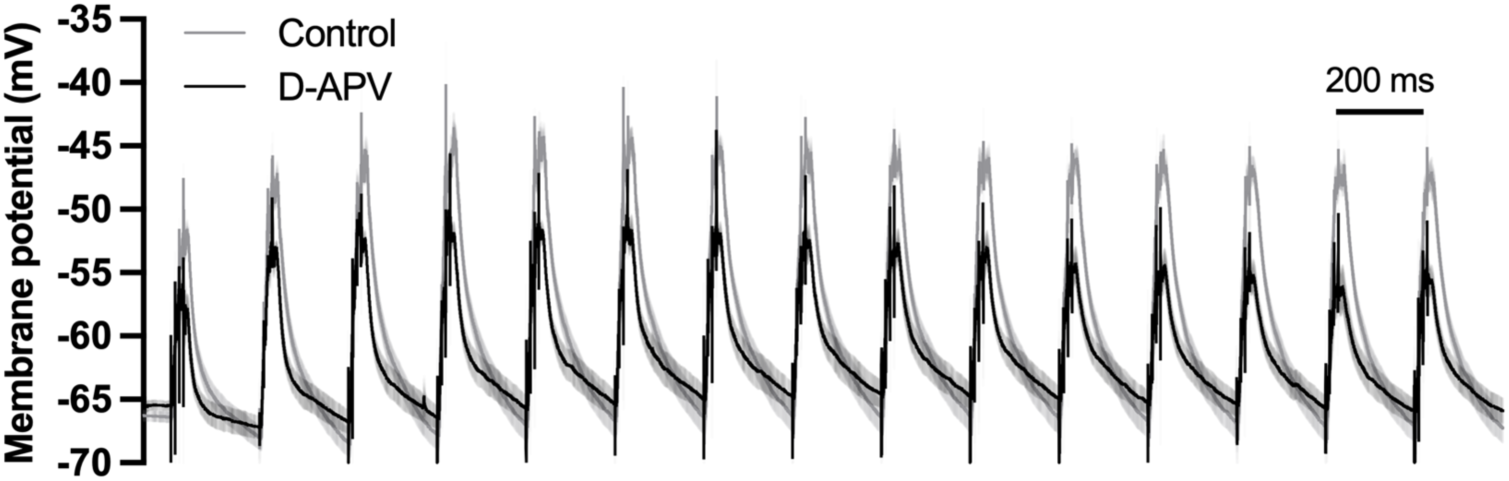
NMDAR-dependent component of synaptic potentials during bursting activity. PSPs during TBS from CA1 pyramidal neurons in hippocampal slices treated with D-APV (50 µM) (n = 8) or untreated controls (n = 15) combined from GluN1a and GluN1b mice (Figure 5D). Action potentials were truncated to examine the underlying PSPs. Data are represented as mean ± SEM.

